# MagPEG: a complete extracellular vesicle isolation/analysis solution

**DOI:** 10.1101/2022.10.18.512792

**Authors:** Li Sun, Sara York, Brandon Pate, Yanping Zhang, David G. Meckes

## Abstract

Current extracellular vesicle (EV) isolation methods depend on large expensive equipment like ultracentrifuges and are laborious and time consuming. There is also currently no method available for high throughput isolation to meet clinical demands. Here, we present a method that combines our previous published ExtraPEG method and magnetic beads. Western blot and nanoparticle tracking analysis (NTA) of the purified EVs revealed higher or equivalent recovery and purity with this method compared to ExtraPEG or size exclusion chromatography (SEC) methods. With this newly developed workflow and automated liquid handling instrument, we have successfully isolated up to 96 EV samples from 5 µL pre-cleared serum in 45 minutes. Moreover, DNA / small RNA / protein purification and profiling steps could be seamlessly integrated into the isolation workflow. To profile EV protein markers, EVs were lysed from the binding step and covalently bound to the surface of the beads. TotalSeq or ELISA antibody can be applied with under a standard protocol. With this extended protocol, researchers can easily complete EV isolation and protein profiling experiment within 8 hours. Taken together, we provide a high throughput method for EV isolation and molecular analyses that may be used for sensitive biomarker detection from biological fluids.

## 1. Introduction

Extracellular vesicles (EVs) are defined as membrane vesicles derived from cells, including apoptotic body, microvesicles and exosomes in the order of their size. Exosomes are in the size range from 50 nm to 200 nm and distinguished from other EVs by content and location of their biogenesis [1]. Whereas other extracellular vesicles, like microvesicles, bud off from the cell surface, exosomes are generally regarded as vesicles that originate at internal endosomal membranes of multivesicular bodies (MVB), and are released into the extracellular milieu following fusion of the MVB with the plasma membrane.

Exosomes share several same features to virial particles, especially enveloped virus, like size, cargos and machinery to shed from cells [2]. Exosomes also contain molecular signatures of their progenitor cells, especially carrying viral proteins from infected cells [3]. Adopted from virology field, current widely used exosome isolation methods include ultracentrifugation, precipitation, ultrafiltration, size exclusion chromatography (SEC), ion exchange chromatography [4] etc. In the aspect of high throughput isolation, methods involve ultracentrifugation are definitely incompatible. Exclude this most widely used technique, the candidates left only precipitation-, filtration-, chromatography- and beads-based method. Without combination with other methods, filtration or precipitation could not provide acceptable result on purity. Beads-based antibody pull-down has the highest purity compared with other method, but it’s still time consuming and expensive [5, 6]. Chromatography-based technique are developed also from virus applications, and there is also commercial product of ion exchange membrane in 96 well plate format (Pall, AcroPrep with Mustang Q membrane). Based on our polit experiment of this product, EV purity is low comparing with ExtraPEG and most importantly we failed to find positive result on EV markers (data not shown). Recently, there is a publication using ion exchange beads to isolate EVs [7] which provide another possibility on high throughput experiment. Norgen’s EV isolation kits also use beads slurry to pull down EVs, but may base on pH to facilitate binding and elution. SmartSEC (SBI) provide a combined technique designed for high throughput purpose, which use pored matrix to trap contaminate and SEC to fractionate EVs. For some reason, we did not fully compare these commercial products with our MagPEG. But from the aspect of cost, none of them fit the need of high throughput of clinical screening / diagnostic purpose [8].

Get hints from DNA purification method of polyethylene glycol (PEG) and magnetic beads combination [9], we assumed EV will also be enriched under the similar protocol. Here, we extend our previous ExtraPEG [10] to MagPEG (*<u>Mag</u>netic beads and <u>PEG</u>-based protocol)*. By using this method, EVs were isolated with similar yield, purity and protein markers. We also evaluated our MagPEG protocol’s ability on high throughput screening. Moreover, some original applications of this carboxyl modified magnetic beads had been fully tested and integrated into our basic MagPEG protocol. Branched from the basic MagPEG protocol, EV associate DNA, small RNA and protein could be isolated by using extended protocols. For protein profiling purpose, EV proteins can be conjugated onto the same beads and further quantified by standard beads ELISA or our TotalSeq assay. Overall, we developed a magnetic beads and PEG combined EV isolation method derived from ExtraPEG. The MagPEG protocol is simple, inexpensive, scalable and analysis integrated method to enrich high purity EVs from serum or cell conditioned medium.

## 2. Material & Methods

### 2.1 Cell culture

HEK293 (ATCC, CRL-1573) were cultured in Dulbecco’s Modified Eagle’s Medium (DMEM, Sigma, D5796) supplemented with 10% fetal bovine serum (FBS, ThermoFisher, 26140079) and penicillin-streptomycin (Corning, 30-002-CI). Cells were incubated at 37^°^C in an atmosphere supplemented with 5% CO_2_.

### 2.2 Pre-treatment of cell conditioned medium or human serum sample for EV isolation

EV-depleted FBS was prepared by ultracentrifugation at 100,000□g for 20□hours, then filtered through a 0.2 µm filter. Cells were first seeded in plate with DMEM + 10% FBS until confluent reach 80%. Medium was removed and cells are rinsed with PBS twice. DMEM + 10% EV-depleted FBS are used to continue culture for another 48 hours. Then, harvested medium was cleared by differential centrifuged at 500□g for 5 minutes, 2,000□g for 15 minutes and 10,000 g for 30 minutes to remove cellular debris, apoptotic bodies and large microvesicles.

For MagPEG method, pre-cleared medium was concentrated to about 20-fold by 100KDa cut-off ultrafiltration (Pall, MCP100C41). The spin was carried out as manufacture’s protocol. All the pre-treated medium were immediately used in EV isolation procedure without storage.

De-identified human serum samples were purchased from BioIVT (Hicksville, New York) and stored in −80^°^C. BioIVT declares all samples were collected under IRB approved protocols. Thawed human serum aliquot was diluted 1:1 with particle-free PBS (pass through 0.2 µm filter) and pre-cleared by centrifugation at 10,000 g for 15 minutes to remove large debris. Pre-cleared serum sample was stored in 4^°^C and use within one week.

### 2.3 ExtraPEG protocol for EV isolation

The ExtraPEG method was performed following the previous paper [10].

For cell conditioned medium sample, cleared medium (section 2.2) was mixed with PEG-6000 (Alfa Aesar, AAA17541-0I) stock solution (working conc. PEG 8%, NaCl 0.5M) and incubated at 4^°^C overnight. On the following day, crude EV were pelleted by centrifuged at 3,000 g for one hour. Then, pellet was resuspended in particle-free PBS and ultracentrifuged at 100,000□g for 70□minutes to wash away contaminating protein and PEG. Purified EV were re-suspended in particle-free PBS and store in −80^°^C until use.

For serum sample, pre-cleared serum (section 2.2) was diluted and mixed with PEG stock solution for 30 minutes. After a 3,000 g centrifugation, the pellet was resuspended in PBS and further purified with 70□minutes ultracentrifugation at 100,000 g. Purified serum EV were resuspend in particle-free PBS and store in −80^°^C until use.

### 2.4 Magnetic beads preparation for MagPEG protocol

SeraMag beads were purchased from Cytiva life science with the stock concentration of 50 µg/µL. Hydrophilic and hydrophobic beads (65152105050250 and 4515210505250) were mixed at equal volume to avoid binding bias. Fifty microgram mixed beads were diluted with PBS into a PCR tube for a single sample isolation protocol. Beads were immobilized on magnet (ThermoFisher, AM10050) and washed three times with PBS. After remove all the buffer, X µL of 50% PEG and Y µL of 5M NaCl (table 1) were added to beads pellet to make a pre-mixed EV binding solution.

**Table 1.** Preparation of pre-mixed EV binding solution

| PEG/NaCl | X µL of | Y µL of | Z µL add to |
| --- | --- | --- | --- |

| Final Conc. in sample | 50% PEG | 5M NaCl | 100 μL sample |
| --- | --- | --- | --- |
| 4% 0.2M | 9.1 | 4.5 | 13.6 |
| 4% 1.0M | 11.1 | 27.7 | 38.8 |
| 6% 0.2M | 14.3 | 4.8 | 19.1 |
| 6% 1.0M | 17.6 | 29.4 | 47.0 |

Pipette up and down slowly several times to resuspend beads until uniform appearance. The amounts of beads, PEG and NaCl were scaled up for multiple or large volume samples. The pre-mixed EV binding solution can be stored in 4^°^C for at least two weeks. Before use, pre-mixed EV binding solution need to be equilibrated to room temperature and fully resuspended with pipette.

### 2.5 MagPEG basic and high throughput protocol for EV isolation

One hundred microliter of concentrated cell conditioned medium or pre-cleared serum sample (from section 2.2) were mixed with Z µL EV binding solution (table 1) in a PCR tube and incubated at room temperature for 10 minutes with agitation. Value Z depends on the final PEG / NaCl concentration needed for each experiment. The tube was spun at 250 g for 2 minutes in a swing bucket centrifuge and placed on magnet for about 2 minutes to immobilize beads. If necessary, slowly pipette the solution up and down to help beads move towards the magnet. After the beads fully migrated, solution was removed. While the tube is on magnet, beads pellet was washed twice with PBS with 30 seconds each time. After removed all the PBS, beads associated with EV can be used for nucleic acid / protein isolation or protein conjugation reaction from this point (section 2.13, 2.15, 2.16).

To eluted EV for NTA or western blot experiments, the tube was removed from magnet and the beads pellet was fully resuspended in at least 20 µL PBS. After incubation for 2 minutes, the tube was placed back on magnet and supernatant was recovered as MagPEG EV sample.

For manual high throughput isolation, pre-mixed EV binding solution (table 1) was added to each well of 96 well V-bottom plate (Corning, 3897) or a clear PCR plate. Orbit mixer at 1000 rpm was suggested to be used in the incubation and elution step. Multichannel pipette was used to add PBS and must not touch beads pellet to avoid sample loss.

Automated high throughput isolation was performed by using SPT Mosquito HV Genomics and Dragonfly Discovery liquid handling platforms at UF ICBR Gene Expression & Genotyping core (RRID:SCR_019145). First, 5 µL of pre-cleared serum sample was transferred to an Eppendorf twin tec 384-well PCR plate using Mosquito HV Genomics system, followed by adding 5 µL of EV binding solution to each well using Dragonfly Discovery. After 30-second mixing with plate shaker and quick spin, the plate was incubated at room temperature for 10 minutes to allow EV binding to beads. Then, plate was placed on 384-well magnet stand for 1 minute and supernatant was transferred to waste plate with two times aspiration (each aspiration is 5 µL). For two time washing, 9.5 µL of PBS is added by Dragonfly Discovery, leave at magnet for 1 minute, then aspirate out to waste plate. Finally, 10 µL PBS was added and the eluted product was transferred to a new 96-well plate for further analysis.

### 2.6 Size exclusion chromatography for EV isolation

SEC column (qEV, Izon, SP1) was used to compare with MagPEG for serum EV isolation [11]. One milliliter of pre-cleared human serum sample was loaded on the column and particle-free PBS was used as mobile phase. EV fraction was enriched in 3 to 6 mL based on refence. All EV fractions were combined and concentrated with ultrafiltration (100KDa cut-off) for characterization and western blot.

### 2.7 EV characterization by Nanoparticle tracking analysis (NTA) and protein quantification

NTA was performed to calculate EV concentration and size by using the NanoSight LM10 (Malvern) with set the camera level to 13 and detection threshold to 5.

Total protein level of EV was quantified by Pierce 660nm Protein Assay (ThermoFisher, 22660) in present of 0.1% SDS to lysis EVs. Protein conjugated with carboxyl beads was quantified by microBCA kit (ThermoFisher, 23235).

### 2.8 Transmission electron microscopy

Sample preparation was similar to previous methods described for exosome electron microscopy [12]. Ten microliter of EV sample were dropped onto Parafilm for each sample preparation. With forceps, a carbon coated 400 Hex Mesh Copper grid (EMS, CF400H-CU-50) was positioned with coating side down on top of each drop for 60 minutes. Grids were then washed with sterile filtered PBS three times, transferring the grid on top of a 30 µL drop of PBS each time. EVs were fixed by positioning the grid on top of a 20 µL drop of 2% paraformaldehyde (EMS, 157-4) for 10 minutes. PBS wash was repeated three times. Post-fixation, grids were then transferred on top of a 20 µL drop of 2.5% glutaraldehyde (EMS, 16537-16) for 10 minutes, followed by three washes with particle-free filtered water. Grid samples were stained on a 20 µL drop of 2% uranyl acetate (EMS, 22400-2) for 10 minutes before embedding on 20 µL of 0.13% methyl cellulose (Alfa Aesar, AA45490-22) and 0.4% uranyl acetate for another 10 minutes. The samples were visualized on an FEI CM120 transmission electron microscope in Biological Science Imaging Resource (BSIR) of FSU.

### 2.9 Western blot

EV samples were lysed by directly mixing with 5X SDS-PAGE sample buffer and boiled. Equal amount of protein from EV lysate were loaded into 10% sodium dodecyl sulfate and polyacrylamide gel (SDS-PAGE). For immunoblotting, proteins were transferred to a nitrocellulose membrane (GE Healthcare, #10600002). The membranes were blocked in a standard TBS-T buffer with 5% non-fat dry milk for one hour at room temperature. Membranes were then probed for EV markers and other proteins with primary antibodies (Table 2) by overnight incubation at 4^°^C. These primary antibodies were subsequently probed with HRP conjugated anti-mouse IgG (GeneTex, GTX213112) or anti-rabbit IgG (GeneTex, GTX77060). ECL substrate was added for picogram (Azure, 10147-296) protein detection thresholds. Chemiluminescence signal was detected using the ChemiDoc MP gel imager (Bio-Rad).

**Table 2.** Antibodies used in the Western blot

| Name | Catalog # |
| --- | --- |
| CD63 | ThermoFisher, 10628D |
| CD81 | GeneTex, GTX72476 |
| CD9 | Abcam, ab2215 |
| TSG101 | GeneTex, GTX70255 |
| EGFR | SantaCruz, sc-03 |
| Syntenin | SantaCruz, sc-100336 |
| HSC70 | SantaCruz, sc-7298 |
| Calnexin | SantaCruz, sc-11397 |
| Alix | SantaCruz, sc-49268 |
| Flotillin-2 | SantaCruz, sc-25507 |
| Vimentin | SantaCruz, sc-6260 |
| Integrin β1 | Cell Signaling, 4706S |
| ApoB | SantaCruz, sc-13538 |
| FN1 | SantaCruz, sc-73611 |
| Albumin | SantaCruz, sc-271605 |
| β-Actin | Cell Signaling, 3700S |
| Histone H4 | Sigma, 07-108 |

### 2.10 Coomassie Blue G-250 staining

After running EV samples on 8-20% SDS-PAGE, gel was removed from the electrophoresis chamber and placed in enough 0.5% Coomassie Blue G-250 (prepared in 50% methanol, 10% acetic acid) to cover the gel. Gel was stained for about 30 to 60 minutes. Stain solution was discarded and gel was rinse briefly with MilliQ water to remove most of the residual stain in the glassware. Destain solution (40% HPLC grade methanol, 10% acetic acid) was added and replaced every 20-30 minutes until faint bands were observed and background was clear. Image was taken by the ChemiDoc MP gel imager (Bio-Rad).

### 2.11 Mass spectrometry and data analysis

EV proteins were purified and digested on S-trap micro column following manufacture’s instruction (Protifi). Briefly, 25 µg of EV samples were lysis with 2X SDS lysis buffer (10% SDS, 100 mM TEAB pH 8.5), reduced with 20 mM dithiothreitol (Promega, V3155, DTT) at 95^°^C for 10 minutes and alkylated with 40 mM iodoacetamide (Amresco, RPN6302V, IAA) at room temperature in dark for 30 minutes. After acidification, samples were bind to S-trap micro column and washed three times with binding/wash buffer. Digestion buffer with 2.5 µg of sequencing grade Trypsin and ProteinaseMax (Promega, V5111, V2071) was loaded on the column and incubated at 37^°^C overnight. On the following day, digested peptides were sequential eluted with 50 mM Tetraethylammonium bromide (TEAB), 0.2% formic acid, 0.2% formic acid in 50% acetonitrile (ThermoFisher, 89871C, MeCN) and pooled.

For testing alternative protein purification and on-beads digestion protocol, SP3 was carried out as described in the literature [13]. Started as the same input, 25 µg of EV samples were lysised with 2X SDS lysis, reduced with 10 mM DTT at 56^°^C for 30 minutes and alkylated with 22.5 mM IAA at room temperature in dark for 30 minutes. After quenched with 10 mM DTT, proteins were bind to beads by adding 100% ethanol to 50% finial concentration and incubated at room temperature for 10 minutes on mixer. Then, beads were washed three times with 80% ethanol. On-beads digestion was using the same digestion buffer as in S-trap protocol. On the following day, the supernatant was transferred into a new tube and peptides were cleaned up by binding to new beads in the present of MeCN (final conc. over 95%). Beads were washed three times with 100% MeCN, and peptides were eluted with 2% DMSO.

The eluted peptides from either method were submitted to FSU college of medicine Translational Science Laboratory to be analyzed on the Thermo Q Exactive HF (High-resolution electrospray tandem mass spectrometer) as previously described [14]. Resulting raw files were searched with Proteome Discoverer 2.4 using SequestHT, Mascot and Amanda as search engines. Scaffold (version 4.10) software was used to validate the protein and peptide identity. Peptide identity was accepted if Scaffold Local FDR algorithm demonstrated a probability greater that 95.0% Likewise, protein identity was accepted if the probability level was greater than 99.0% and contained a minimum of two recognized peptides as previously described.

### 2.12 CD63 direct ELISA

Direct ELISA method was used to show CD63 level in either EV enrichment method. Briefly, samples diluted in 100 µL bicarbonate/carbonate buffer were coated on the MaxiSorp plate (NUNC, 44-2404-21) overnight at 4^°^C. After blocking with PBS-B blocking buffer (1% BSA in PBS, 0.05% tween-20) at room temperature for one hour, CD63 antibody (same as in western blot) were added to each well at 1:250 and incubated overnight again to maximize performance. Anti-mouse IgG HRP was used at 1:1000 concentration as secondary antibody and incubated at room temperature for 30 minutes. Between each step, the plate was washed 5 times with at least 300 µL PBS-T (0.05% tween-20 in PBS) to remove any unbonded antibody. One hundred microliters TMB substrate solution (BioLegend, 421101) was developed for 10 minutes in dark or until reach the desired color. After stop with 50 µL 2N sulfuric acid, signals were read at 450 nm.

### 2.13 Nucleic acid isolation after MagPEG

Started from the beads associated with EV step in basic MagPEG protocol (section 2.5), the tube was removed from magnet and beads were fully resuspended with lysis buffer (Tris-EDTA, 0.5% Triton X-100). Incubation could be carried out at room temperature or boiled for 10 minutes, if necessary. Two volume of DNA binding solution (20% PEG, 2.5M NaCl in ddH_2_O) was mixed with EV lysate and beads mixture for another 10 minutes. Beads were immobilized by centrifugation and placed on magnet. While the tube was kept on magnet, beads were washed twice with 70% ethanol and air dried. Purified DNA was eluted with ddH_2_O or TE buffer.

For small RNA isolation, beads were incubated with lysis/protein digestion buffer (Tris-EDTA, 0.5% Triton X-100, proteinase K) at 37^°^C for 30 minutes to fully release small RNAs. Then, 50% PEG and 5M NaCl stock solution were added to mixture to a final concentration of 20% and 1.25M. After incubate at room temperature for 5 minutes, beads bonded with large RNA/DNA molecules were discarded and supernatant was transferred into a new tube. Fresh beads were mixed and discard again to further clarify residual RNA/DNA. After this step, cleared supernatant was mixed with new beads and 3 volume of 100% ethanol to facilitate small RNA binding to beads. Beads were washed with 70% ethanol and air dried. Purified small RNAs were eluted with ddH_2_O or TE buffer.

DNA or small RNA isolated with beads were quantified by Qubit dsDNA or small RNA quantification kit, respectively (ThermoFisher, Q32851, Q32880). Isolated cfDNA and MagPEG EV DNA were analyzed by High Sensitivity DNA Chip (Agilent, 5067-4626) on Agilent 2100 Bioanalyzer.

### 2.14 Cell free DNA (cfDNA) isolation from serum

cfDNA was isolated by magnetic bead-based protocol similar to some commercial kits and widely used protocols [9, 15-18]. Briefly, 100 µL pre-cleared serum sample were mixed with 5 µL Triton X-100 (10%) and boiled for 10 minutes After cooling to room temperature, two volume of DNA binding solution (20% PEG, 2.5M NaCl in ddH_2_O) and 50 µg beads was mixed with lysised sample for another 10 minutes. Beads were immobilized by centrifugation and placed on magnet. While the tube was kept on magnet, beads were washed twice with 70% ethanol and air dried. Purified DNA was eluted with ddH_2_O or TE buffer.

### 2.15 Protein isolation after MagPEG

EV protein was isolated following SP3 protocol as described in the literature [13]. From the beads associated with EV step (section 2.5), the tube was removed from magnet and beads were fully resuspended with lysis / DNA digestion buffer (Qiagen, 79254; buffer RDD with DNase and Triton-X100 (final conc. 0.5%)). The mixture was incubated at room temperature for at least 15 minutes to eliminate DNA contamination. One volume of 100% ethanol was mixed with EV lysate and beads for another 10 minutes to facilitate protein binding with beads. Beads were immobilized by centrifugation, and supernatant was removed while the tube was on magnet. Taking the tube off the magnet, bead pellet was washed twice by fully resuspending the beads in 80% ethanol. Protein was eluted with PBS + 0.5% SDS. Sonication, boil or vortex may be used if the beads were not resuspended well in the elution step.

### 2.16 EV protein conjugation on Beads

Protein conjugation protocol was following the manufacture’s technical application note (Cytiva Procedure 29174976 AA) with modification.

Before the experiment, the amount of N-(3-Dimethylaminopropyl)-N_′_-ethylcarbodiimide hydrochloride (Sigma, 03449, EDAC) used in the reaction was calculated with the following equation:

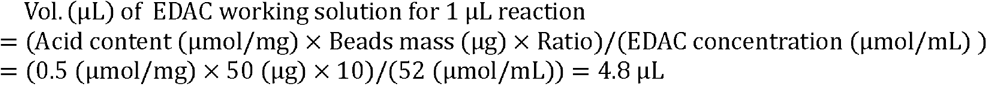

Carboxylic acid content of beads is provided in mEq/g, which is equivalent to µmol/mg. From the test lot, the value is 0.5 mEq/g. For all the protocols mentioned above, 50 µg beads were used in a single sample isolation. EDAC to acid ratio could be adjusted from 1 to 10. In here, we use 10 to maximize protein conjugation efficiency based on the preliminary results (supplementary Table 2). EDAC concentration of freshly prepared solution is 52 µmol/mL.

From the step in section 2.5, beads were resuspended with 20 µL of 4-Morpholineethanesulfonic acid buffer (Sigma, M3885, MES, 50 mM) and incubated on mixer at room temperature for 15 minutes. One hundred microgram of BSA was mixed with beads as a separate control reaction for downstream analysis. Bond to beads were promoted under 50% ethanol as in SP3 protocol. Then, unbonded BSA was removed and beads were resuspended in 20 µL of MES buffer (50 mM) as in other EV samples. Based on the calculation above, 96 µL of freshly prepared EDAC was added to each reaction tube and mixed for one hour on mixer at room temperature. After the reaction was finished, supernatant was removed and beads with coupled EV protein were washed three times with PBS-T. Then, the beads were resuspended in PBS for protein quantification. The protein conjugated beads can be stored in 4^°^C at least for a week before used in beads ELISA or TotalSeq assay for protein profiling. Beads were also run on BD FACSCanto flow cytometry in FSU college of medicine core facility for aggregation test.

### 2.17 Direct Beads ELISA

After protein quantification, 5 µg protein of beads were blocked with PBS-B blocking buffer at room temperature for 30 minutes on orbit shaker (1000 rpm). The CD63, CD9 or CD81 antibody (same as in western blot) was added at concentration of 1:250 dilution in PBS-B blocking buffer and incubated on shaker for two hours. Then, beads were washed 3 times with PBS-T and incubated with HRP conjugated anti-mouse IgG for another 30 minutes. After wash, beads were resuspended with 100 µL TMB substrate solution (BioLegend, 421101) until reach the desired color. After stop with 50 µL 2N sulfuric acid, solutions without beads were transferred into a new plate and read at 450 nm.

### 2.18 TotalSeq assay on beads

TotalSeq assay was performed following the previous paper with modification [19].

The equal protein mass of beads was blocked with casein blocking buffer (2.5% casein in PBS, 0.05% tween-20, 100 µg/mL sheared salmon sperm ssDNA (BioVision, B1677)) at room temperature for 30 minutes on orbit shaker (1000 rpm). A TotalSeq-A antibody pool (TotalSeq-A0404 anti-human CD63 Antibody (353035); TotalSeq-A0373 anti-human CD81 (TAPA-1) Antibody (349521); TotalSeq-A0090 Mouse IgG1, κ isotype Ctrl Antibody (400199); TotalSeq-A0579 anti-human CD9 Antibody (312119)) was added at concentration of 1:2000 dilution in 100 µL casein blocking buffer and the sample was incubated on shaker for two hours. Then, beads were washed 5 times with PBS-T (0.05% tween-20 in PBS). Klenow polymerase (NEB, M0212), dNTP (NEB, N0447) and synthesized 3’-adaptor (see Table 3) were mixed on ice as follow for the extension mix.

| | Vol. ($\mu\text{L}$) | Final Conc. |
| --- | --- | --- |
| 10X Buffer 2 | 4.0 | 1X |
| dNTP (400 $\mu\text{M}$) | 2.0 | 20 $\mu\text{M}$ |
| Klenow | 0.8 | 1 U |
| 3'-Adaptor (10 $\mu\text{M}$) | 2.0 | 500 nM |
| PCR grade $\text{H}_2\text{O}$ | 31.2 | |
|  | 40.0 |  |

**Table 3.**
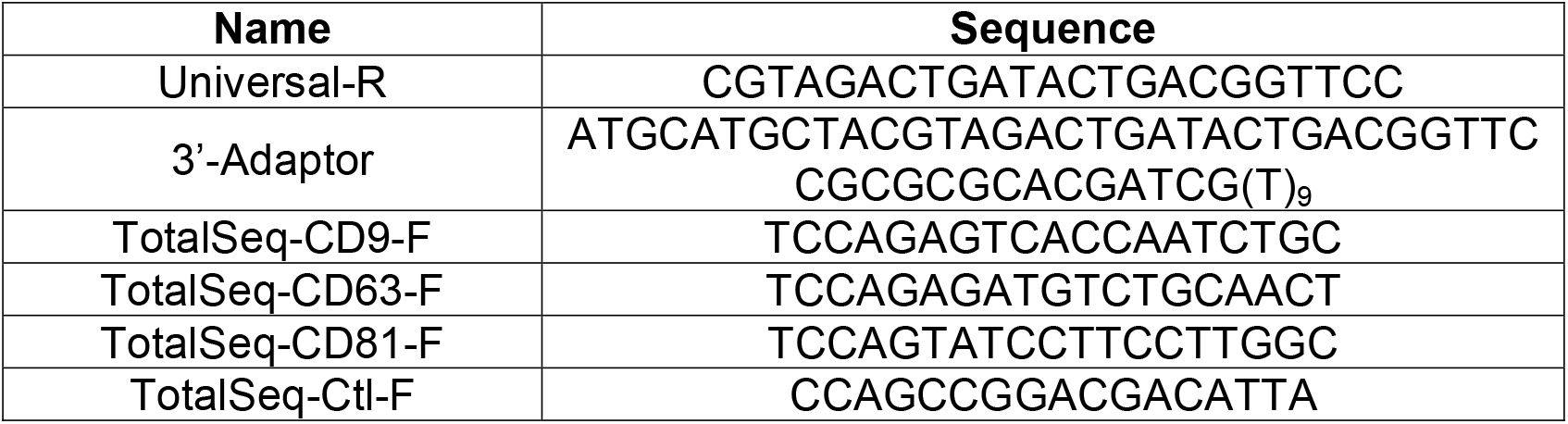
Sequence of TotalSeq Primers and Oligos

The beads were resuspended with 40 µL extension mix and incubated at 37^°^C on a PCR cycler (Eppendorf, UX-93944-39) for 10 minutes. After heat-inactivated at 95^°^C for 5 minutes, beads were cooled down to room temperature and clear supernatant was separated by magnet. One microliter of the supernatant was used as template in 15 µL qPCR reaction containing 2X PerfeCTa SYBR Green FastMix (QuantaBio, 95072), gene specific forward primer and universal reverse primer (see Table 3, 250 nM final conc.). The protocol for qPCR is 95^°^C for 5 minutes, 40 cycles of 95^°^C for 10 seconds, 60^°^C for 10 seconds, 72^°^C for 20 seconds, followed by a melt curve assay to prove specific product in the reaction. The qPCR was carried out on CFX Connect Real-Time PCR Detection System (Bio-Rad) and data were analyzed by CFX Manager Software (Bio-Rad).

### 2.19 Statistical analysis and software

Raw data were collected from the software used with instrument of plate reader or gel imager. Experiments were carried out at least in two to three biological replicates, and data is represented as a mean + standard deviation. Statistical significance was determined by student-t test with p<0.05. Excel software were used for column and scatter chart. Western blot image was chopped to show clear bands, and brightness/contrast was adjusted only if necessary. PCA plot was made by NetworkAnalyst 3.0 [20]. Venn diagram was made by BioVenn [21]. Other diagrams were created by PowerPoint.

## 3. Results

### 3.1 MagPEG combines PEG precipitation and magnetic beads for EV isolation

From the hint of DNA purification protocol by using magnetic beads [9], we reasonably hypothesized that in the present of PEG, EV will also aggregate around the magnetic beads which will be pulled down by magnet in the following step. In the DNA purification protocol, DNA will bond with beads only in the present of PEG and NaCl, which means it’s not a strong interaction with beads [9]. So, we designed the new EV isolation workflow by replacing the wash solution (70% ethanol) and elution buffer (TE) with PBS (Fig.1a). Similar to DNA protocol, wash step will be only preformed when the tube is on magnet. After wash twice with PBS, the tube was removed from magnet and beads were fully resuspended with PBS to elute bond EVs. No particles were eluted from the beads only or beads plus BSA sample at this step (Supplementary Table 1). The protein still binding on the beads after elution step was striped by 0.5% SDS and boiled for testing (Fig.1b-c). From western blot results of these wash, elution and stripping fractions, only the elute enriched EV marker of CD63 and TSG101 (Fig.1b). And protein quantification indicate that wash step removed more than 90% of the contaminate protein without any marker present (Fig.1c). These results prove that our new protocol could purify EV with a protocol similar to DNA purification. And because it was developed from our previous ExtraPEG protocol and combined magnetic beads, so we decided to name it as MagPEG.

**Fig1.**
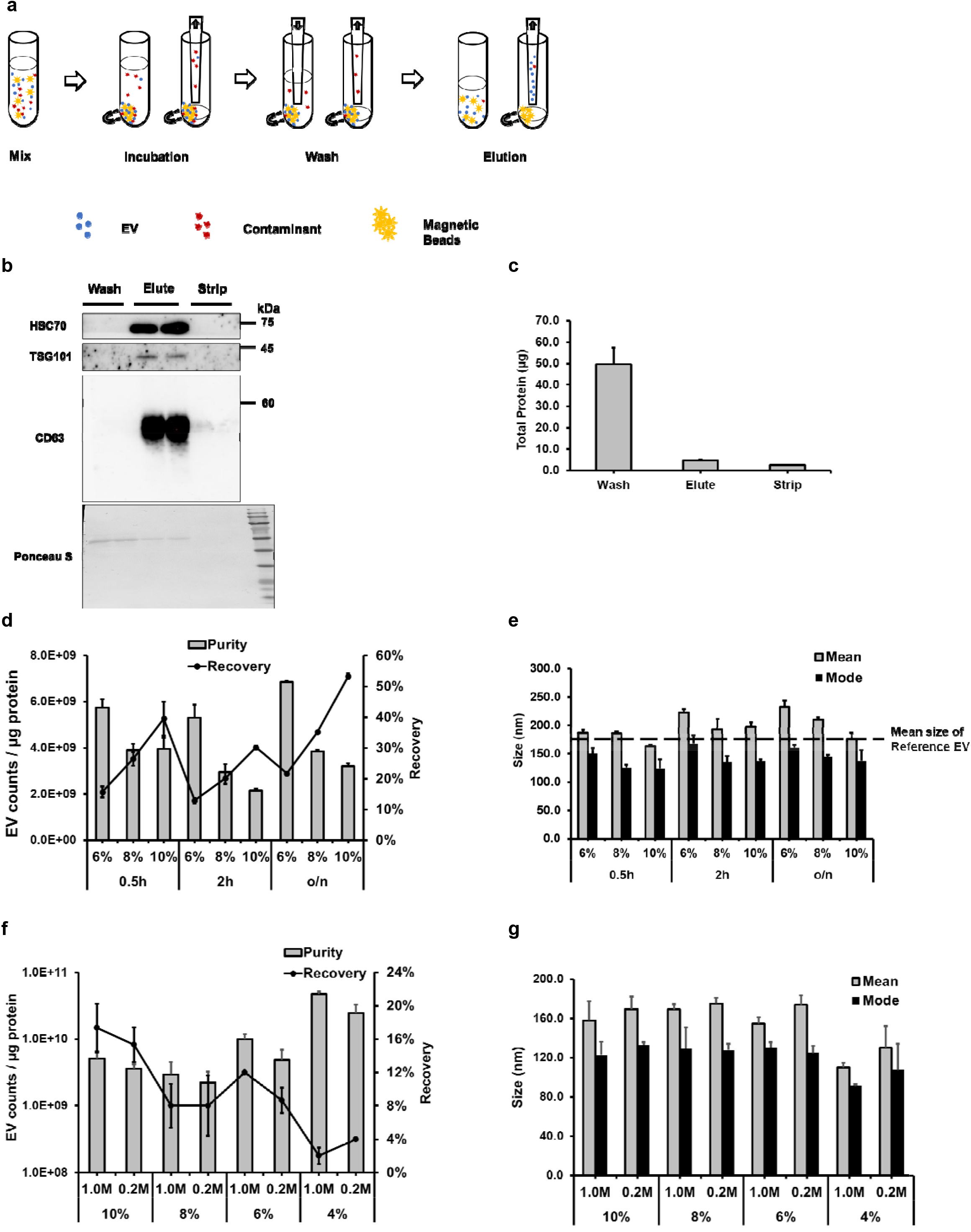
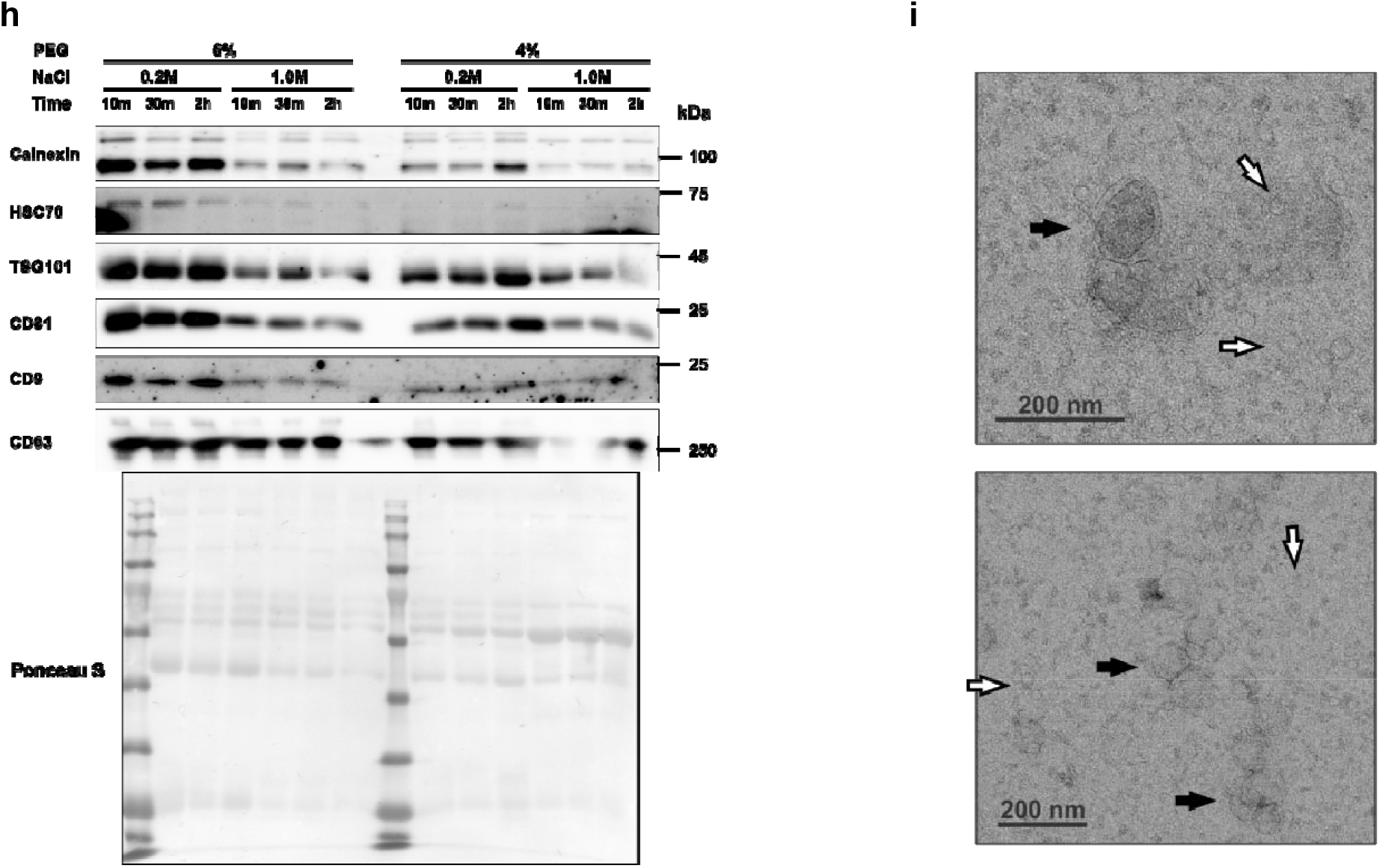
MagPEG Protocol optimization for EV isolation. **(a)** Workflow for MagPEG basic protocol. **(b)** Western blot for EV marker of different fractions from MagPEG protocol. Samples were loaded as equal protein amount. **(c)** Total protein quantification of different fractions from MagPEG protocol. **(d)** PEG concentration and incubation time effect on MagPEG EV’s purity and recovery. **(e)** PEG concentration and incubation time effect on MagPEG EV’s size. Dot line shows the size of input EV sample. **(f)** PEG and NaCl concentration effect on MagPEG EV’s purity and recovery. **(g)** PEG and NaCl concentration effect on MagPEG EV’s size. **(h)** Western blot for EV markers of purified EV from different conditions. Samples were loaded as equal protein amount. **(i)** Electron microscopy image of MagPEG 6% / 0.2M sample. Black arrow shows the classic cup shape vesicles. White arrow shows the small single layer vesicles.

Next step, we tuned different conditions based on EV isolation results like purity, recovery and protein markers. The critical variables in the MagPEG protocol include PEG concentration, NaCl concentration and incubation time. Our pilot study started from a simplified model, EV-depleted medium spiked in with purified EV which will provide an excellent reference. Similar to ExtraPEG result, higher PEG concentration (8% and 10%) will increase yield but lower the purity (Fig.1d). Longer incubation time have the similar effect on yield and purity (Fig.1d). But in the 6% group, longer than 2 hours significantly increase size of purified EV (Fig.1e). So, on the second-round experiments, we focus on lower PEG concentration (4%, 6%), shorter incubation time (10 minutes, 30minutes), NaCl concentration (0.2M, 1.0M) and comparison with ExtraPEG method. High salt concentration trend to increase the purity but not significate (Fig.1f). For the PEG concentration from 10% to 6% despite of NaCl, the EV size was all in typical exosome range (100 - 200nm), but 6% / 1.0 M combination gave the highest purity results (Fig.1f-g). We also test different type of beads (hydrophilic and hydrophobic) on MagPEG EV isolation (Supplementary Fig.1). The results showed that change the PEG concentration had greater impact than beads type. The only interesting result is hydrophobic beads enriched more CD81+ EVs than hydrophilic beads under 6% PEG. So, for the standard MagPEG protocol, we always mix the hydrophilic and hydrophobic beads in 1:1 ratio as other researcher did in SP3 protocol [13].

To complete research all conditions in one experiment, we used pre-cleared serum as input and probed EV markers on those EV isolated from different conditions. From the western blot result (Fig.1h), 6% PEG / 0.2M NaCl group had the highest CD63, CD9, CD81 and TSG101 level among all the groups. There is no difference within each group of different incubation time, which confirm that 10 minutes incubation is sufficient for EV isolation. The most surprising results were coming from the 4% PEG / 1.0M NaCl combination, which has the smallest size (mode < 100 nm), highest purity (> 5.0 E10 particle / µg protein) and dimmest EV marker (Fig.1f-h). As we were using the serum sample, one of the reasonable guesses is this PEG / NaCl combination enriches HDL or LDL particles more than exosomes. Some previous studies were also using PEG to isolate LDL particles [22, 23]. EM images of 6% / 0.2M sample gave us the similar impression. Not only the classic cup shape vesicle (100 – 200 nm, black arrow in Fig.1i) were found in this sample, but also a lot of smaller single membrane vesicles (<50 nm, white arrow in Fig.1i) present in the same image (Fig.1i).

Concluded all these results, we chose to use PEG 6%, NaCl 0.2 M and 10 minutes incubation time for further experiment to enrich most wide EV subpopulations for study.

### 3.2 MagPEG enriched EV from serum and concentrated conditioned medium sample

To fully characterize of MagPEG purified EV, we used serum and cell conditioned medium as input to test. On the serum sample, we also compared with previous ExtraPEG method and widely used SEC method [11]. Compared with ExtraPEG, MagPEG have similar size distribution and purity, but significantly higher recovery (Fig.2a-c). SEC purified smaller EV with less protein, and comparable recovery as MagPEG (Fig.2a-c). From the western blot, MagPEG had comparable level of CD63 and TSG101, but lower CD9 and CD81 than ExtraPEG (Fig.2d). For other makers, MagPEG enriched low molecular weight EGFR (Fig.2d). Almost on all the proteins tested by western blot, SEC present the faintest bands in the three methods. Coomassie Brilliant Blue (CBB) and ponceau S staining also showed that different protein patterns of enriched EVs by those methods (Fig.2d-e). The clearest result from the CBB staining is MagPEG had significantly less albumin (67 kDa) accumulate which may due to shorter incubation time. And MagPEG also had different bands on large molecular weight region (180 kDa and >250 kDa) which may come from LDL’s APOB as we assumed in the pilot study (Fig.1i).

**Fig2.**
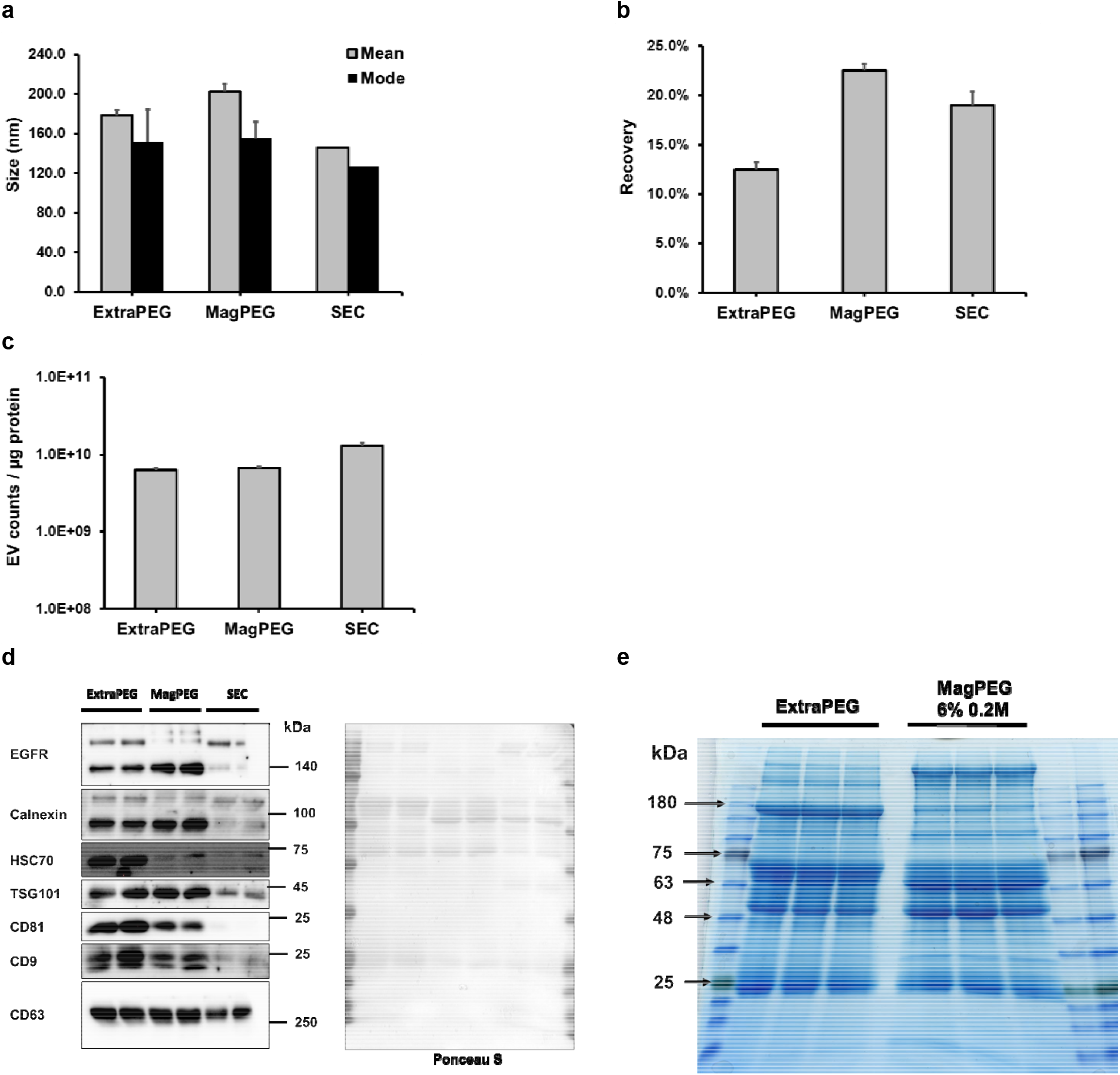
Characterization of MagPEG purified EV from serum sample. **(a)** EV size from different isolation methods. **(b)** EV recovery from different isolation methods. **(c)** EV purity from different isolation methods. **(d)** Western blot for EV marker from different isolation methods. Ponceau S staining shows the equal protein loading for each lane. Sample were run in duplicate and equal protein amount. **(e)** Coomassie Bright Blue staining of EV isolated from ExtraPEG and MagPEG 6% / 0.2M. Samples were run in triplicate.

When we test MagPEG on cell culture medium, the results were a little different comparing with serum sample. EV concentration in cell culture medium is relative 2 magnitude lower than serum (10^7^ vs. 10^9^ per µL) [24]. Low EV concentration requires longer incubation time or higher PEG concentration to archive acceptable recovery, but also aggregate more contaminate proteins which lower the purity (Fig1d-e). Unlike ExtraPEG, on-magnet beads washing step may not efficiently remove lipoproteins as ultracentrifugation. The only way to resolve the dilemma is to increase the EV concentration before apply MagPEG. So, ultrafiltration (100kDa cut-off) was used to concentrate cell culture medium. Due to the limitation of ultrafiltration device, 20-fold enrichment is the highest concentration factor we could get. Using this concentrated sample, we preformed same condition (6%, 0.2M, 10 minutes) of MagPEG protocol as in serum. Comparing with ExtraPEG, EV had similar size, purity and recovery (Fig.3a-c). But western blot showed that MagPEG EV from concentrated medium had lower level of some exosomal maker proteins like TSG101, CD81, Flotellin, Alix, but similar density of others like CD9, CD63, Syntenin and Vimentin (Fig.3d). These results prove again that MagPEG enriched different subpopulation of EVs comparing with ExtraPEG. There were also difference between ExtraPEG and MagPEG on some non-EV marker proteins like Integrin β1 and EGFR. To evaluate contaminate level, extend exposure (about 5 minutes) of Calnexin was used to visualize clear band. ExtraPEG perform better in serum sample, but not in medium sample (Fig.2d, Fig.3d). Another result needs to be notice is that HSC70 is almost absent in all the western blot results of MagPEG EVs regardless of sample type or conditions (Fig.1h, Fig.2d, Fig.3d). HSC70 has been widely considered as an EV marker which is present in the EVs isolated from all the other methods [25, 26]. But our results proved that HSC70 is not a crucial marker for EV or exosome, and it may belong to an unknown EV subpopulation or protein aggregates.

**Fig3.**
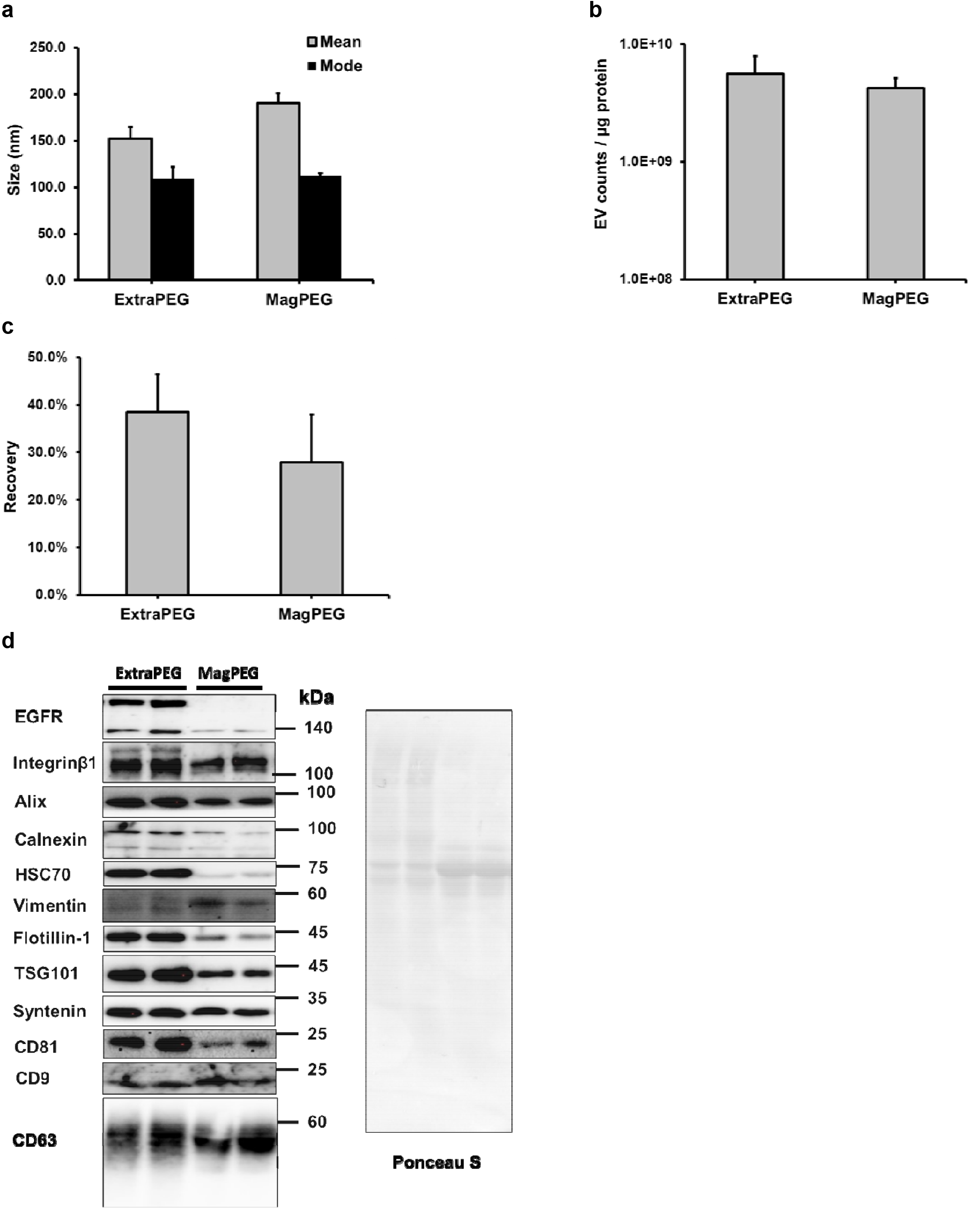
Characterization of MagPEG purified EV from cell conditioned medium. **(a)** EV size from different isolation methods. **(b)** EV purity from different isolation methods. **(c)** EV recovery from different isolation methods. **(d)** Western blot for EV markers from different isolation methods. Ponceau S staining shows the equal protein loading for each lane. Samples were run in duplicate.

### 3.3 Mass spectrometry analysis reveal MagPEG enriched different subtypes of EV

To visualize protein composition of MagPEG, we purified EV proteins from ExtraPEG or MagPEG with four different PEG / NaCl combinations (6%/0.2M, 6%/1.0M, 4%/0.2M, 4%/1.0M). S-trap micro column was used to remove PEG residue in sample. The mass spectrometry identified total 427 unique proteins in all groups (supplemental file). As shown in PCA plot, biological triplicates from each group were well clustered together (Fig.4a). Protein list from two of MagPEG groups (6% / 0.2M and 4% / 1.0M) were compared with ExtraPEG (Fig.4b). About 214 unique proteins (64% of total 333) were present in all three groups, and each group also had its signature proteins listed in Table 4. This heterogeneity was present not only on the identified proteins, but also the quantification results when comparing MagPEG (6% / 0.2M) and ExtraPEG (R^2^ = 0.44, Fig.4c). Some of protein quantification results from MS were shown in Fig4d. The wildly identified cell plasma reference protein GAPDH and ACTB were very low or undetectable in the group of 6%/0.2M and 4%/1.0M which may indicate that these conditions enrich less contaminate cytoplasmic proteins. This same result on the histone protein H4 may also indicate less genomic DNA contamination in the MagPEG EVs. Moreover, for some extracellular soluble proteins like albumin (ALB) or transferrin (TF), 6%/0.2M is the lowest among all the groups. The most important discovery of proteomic is MagPEG did enrich LDL particles because the marker protein APOB is significantly higher in all MagPEG groups comparing with ExtraPEG sample. And two conditions (6%/1.0M, 4%/0.2M) also had LDLR in their protein lists. Another prove of these highly heterogenous results were came from transmembrane receptors like CD14 (monocytes marker) and ITGA2B.

**Fig4.**
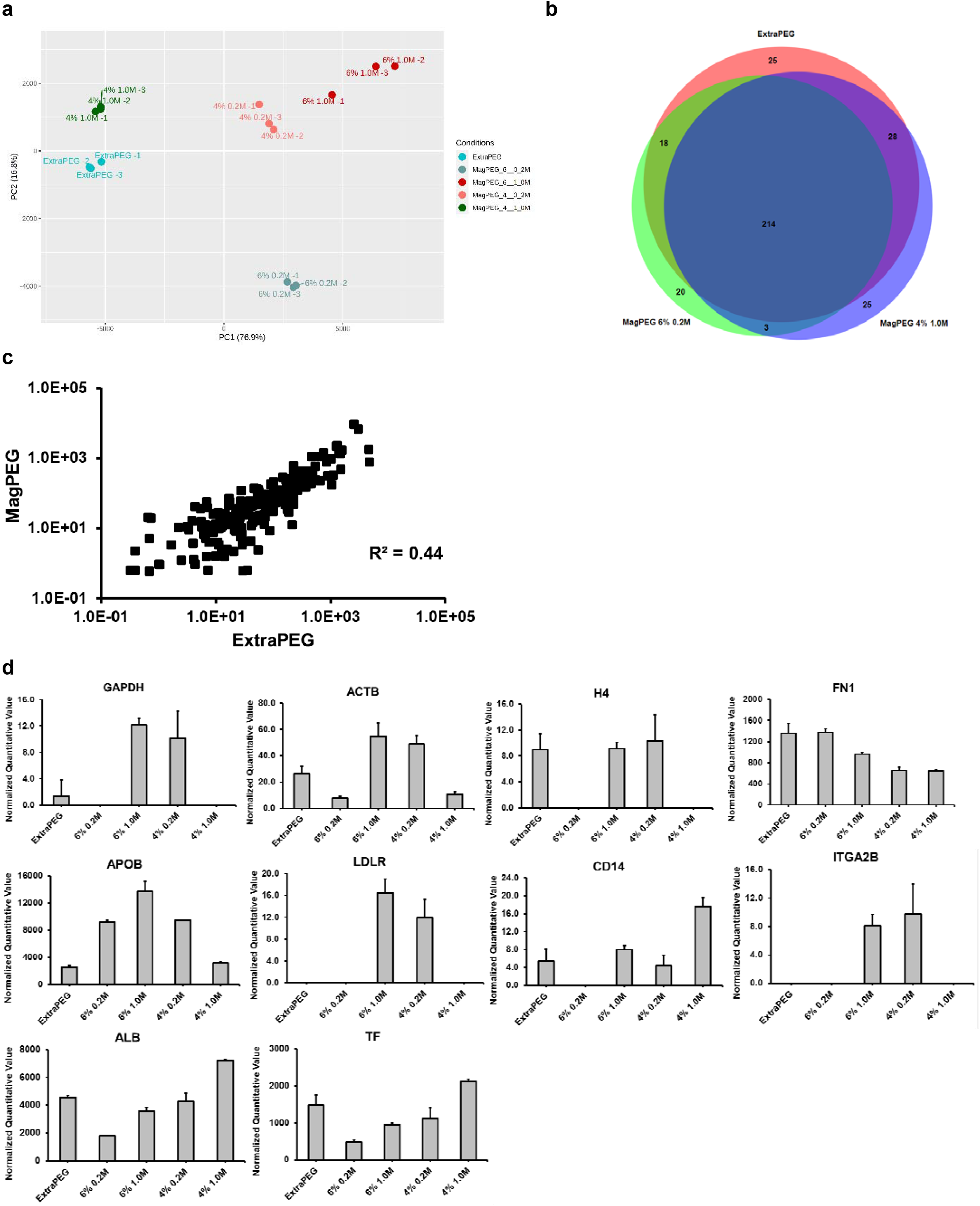
Proteomic analysis of MagPEG purified EVs. **(a)** PCA plot of each sample group. Samples in each group were run in biological triplicate. **(b)** Venn diagram of unique proteins in ExtraPEG, 6% 0.2M and 6% 1.0M group. **(c)** Quantitative value of identified protein from ExtraPEG and MagPEG 6% 0.2M group. **(d)** Normalized quantitative value of selected proteins in all the groups.

**Table 4.** List of unique proteins present only in the groups

| ExtraPEG only | 6% 0.2M only | 4% 1.0M only |
| --- | --- | --- |
| LRP1 | PLA2G7 | BTD |
| IGKV1-5 | COMP | CST3 |
| H4-16 | APOF | SELL |
| FTL | IGHV3-48 | SFTPB |
| H2BC9 | IGHV3-43 | OTOF |
| RAP1B | MARCO | HNRNPM |
| S100A8 | PCSK9 | VASN |
| S100A9 | BPIFB1 | TBC1D2 |
| DSG1 | SLIT1 | CHMP4A |
| UBA52 | SPTBN4 | INPP5D |
| GAPDH | FBN1 | LCP1 |
| H2AC18 | WASHC2A | HSPA7 |
| IGLV5-37 | IGKV1D-13 | LYVE1 |
| KIF5C | TNC | ABCA1 |
| CEP170 | BSN | TLN2 |
| DEFA3 | THBS1 | SPP2 |
| ANPEP | SHANK1 | PTGDS |
| FCN1 | CASP14 | AKAP11 |
| F7 | ZC3H11A | CLVS2 |
| INTS4 | HYDIN | FRMPD1 |
| SACS |  | MAN1A1 |
| LRRC71 |  | AKAP9 |
| PCLO |  | COL4A2 |
| CRISP3 |  | KMT2D |
| GVINP1 |  | TP53BP1 |

Western blot was used to verify Mass spectrometry results (Fig.5). APOB and FN1 had similar results in these two different protein profiling methods. Some of the EV markers also repeat our pervious results as shown in Fig1h and Fig.2d. The 6% / 0.2M group and ExtraPEG had the lowest albumin and HSC70 level, and comparable quantity on EV markers like TSG101, CD81, CD9. The only difference is APOB which imply MagPEG 6% / 0.2M not only enrich the same population as ExtraPEG, but also LDLs. MagPEG 4% / 0.2M also had similar EV marker panel with ExtraPEG, but with less LDLs and more other contaminate proteins (albumin and HSC70). This result provides us an option that PEG concentration can be switched between 6% and 4% to meet the need of research goal for EV subpopulations. The 4% / 1.0M did not enrich LDLs as we expected from Fig.1f-h. The high particle concentration and small size may come from protein aggregates of albumin (or HSC70). From the MS and western blot results shown here, we could confirm that MagPEG protocol has ability to purify distinct EV subpopulations under different conditions.

**Fig5.**
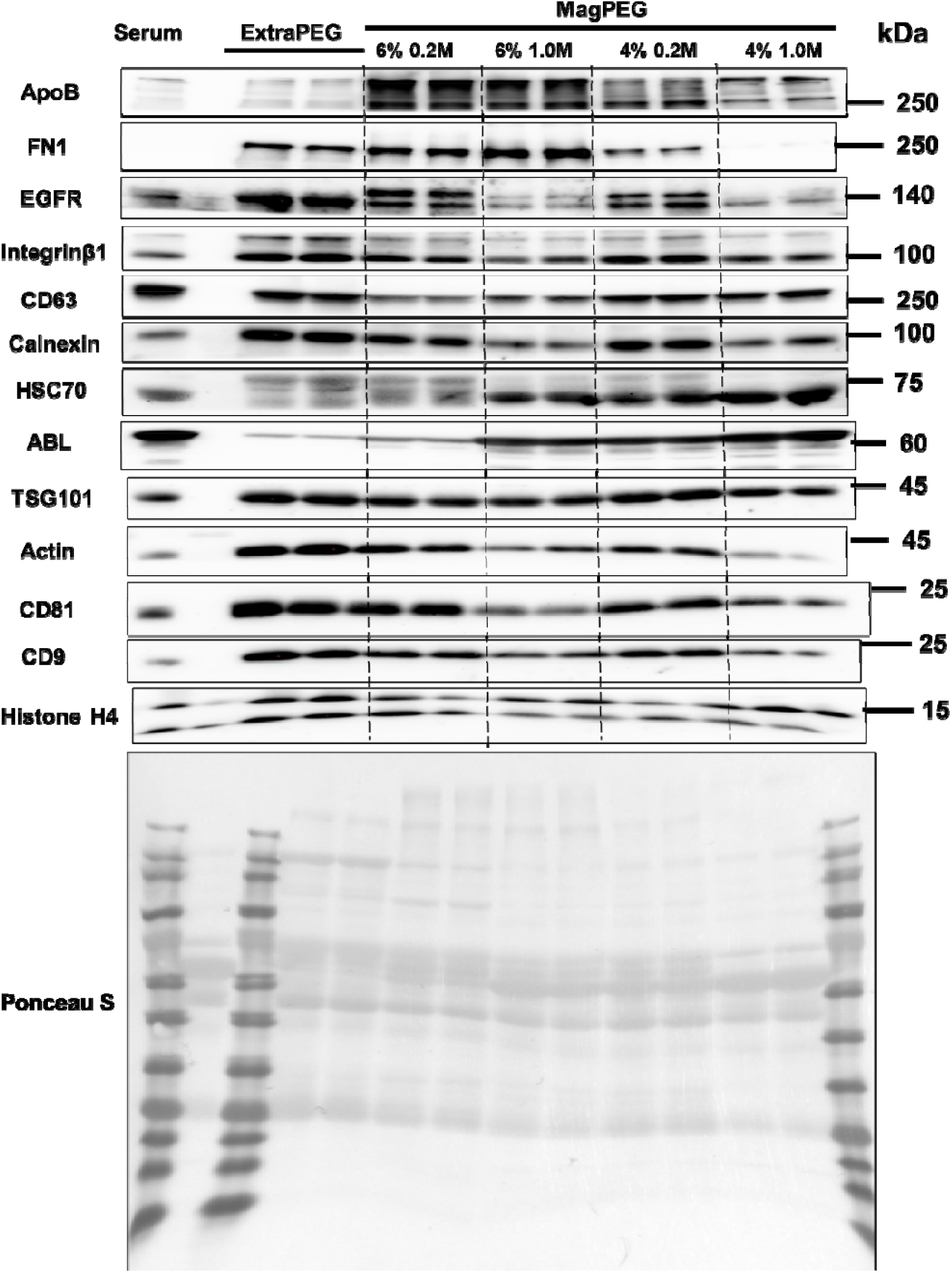
Western blot analysis of ExtraPEG and MagPEG EVs. Biological duplicate samples were run in each condition. Ponceau S staining shows the equal protein loading for each lane.

Another interesting application of this magnetic beads is to purify protein and peptides for mass spectrometry experiments, which is also known as SP3 protocol [13]. In here, we used the same MagPEG 6% / 0.2M sample as input, and EV proteins were purified with either S-trap column or SP3 protocol. The proteins identified by MS were quite similar (Fig.6a) and the quantification were also highly corelated (Fig.6b, R^2^ = 0.96) between these two methods. Here, we repeat the results from previous literature [13], and expand its application on EV proteins.

**Fig6.**
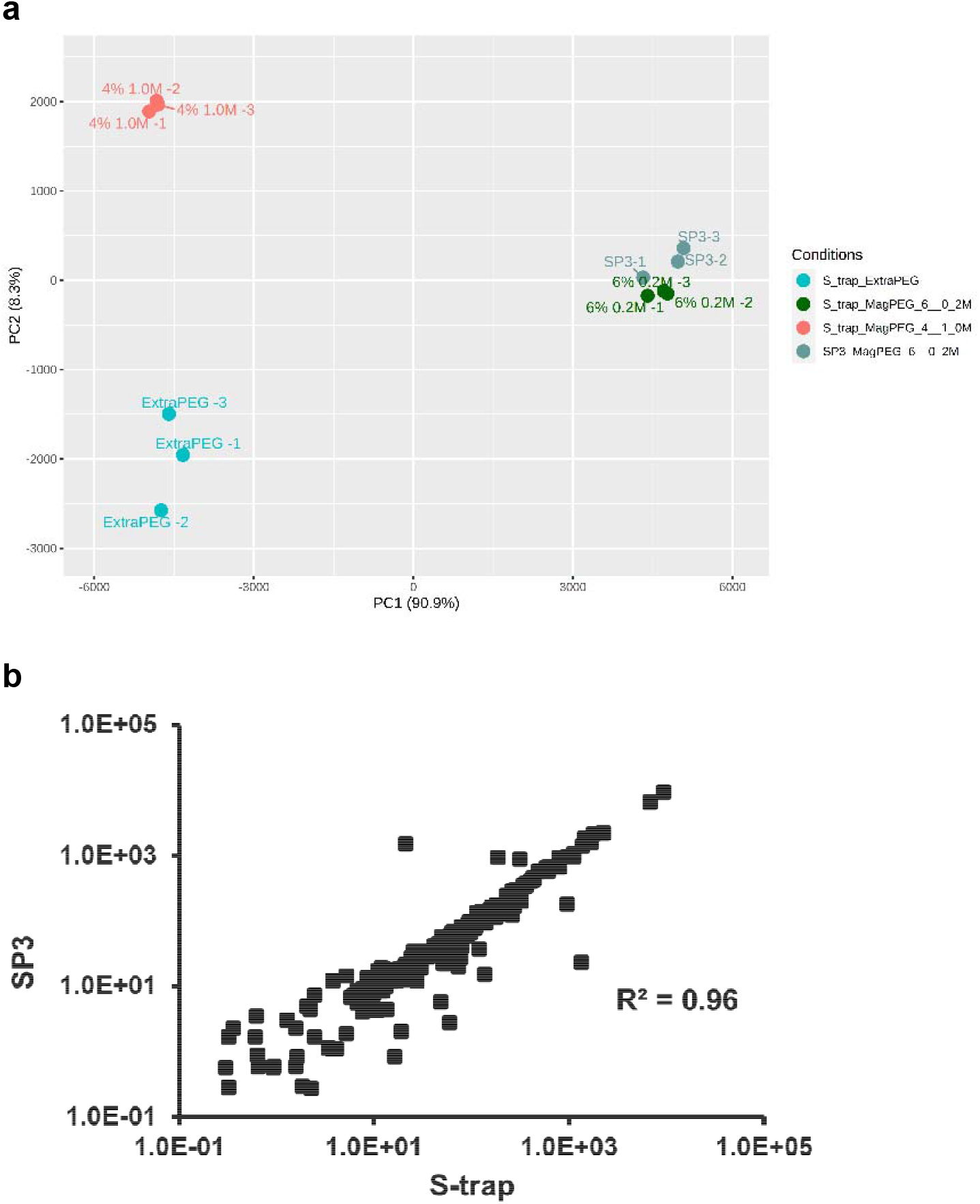
Comparison of S-trap and SP3 method for proteomic sample preparation. **(a)** PCA plot of each sample group. Samples were run in triplicates. S-trap purified ExtraPEG samples and MagPEG 4% 1.0M samples were shown as external reference. **(b)** Quantitative value of identified protein from S-trap and SP3 method.

### 3.4 MagPEG protocol has great potential on high throughput EV isolation

The basic MagPEG protocol used only magnetic beads, magnet, common lab supplies and chemicals, which provide an excellent platform to expand its capacity without increase the cost. So, we focused on regular lab equipment, like multichannel pipette and microplate, to test the high throughput ability of MagPEG. Almost the same step as basic protocol, pre-mixed EV binding solution were added into a clear v-bottom plate, and mixed with sample on an orbit shaker to archive consistent result between wells. Low speed centrifugation and clear v-bottom plate will make sure the visibility of beads during the whole procedure. By using this modified manual high throughput protocol, we had successfully purified up to 32 serum EV samples within 30 minutes. From 50 µL pre-cleared serum, we purified EV samples with about 20 µg total protein, all of them had a protein concentration around 1.0 µg/µL (Fig.7a). Most importantly, all the samples had detectable EV marker CD63 which corelated with their protein concentration (Fig.7b-c, R^2^ = 0.55). We also randomly chose 12 samples and use equal volume to run western blot for more EV makers as shown in Fig7d. All of the markers (CD81, CD63, TSG101) shows consistent results but also variations. Sample in lane 2 had the highest of EV markers with lowest HSC70, and lane 1 is the dimmest of all. Notice that HSC70 is under a extend exposure to visualize the band as previous Fig.1h, Fig.2d, Fig.3d and Fig.5.

**Fig7.**
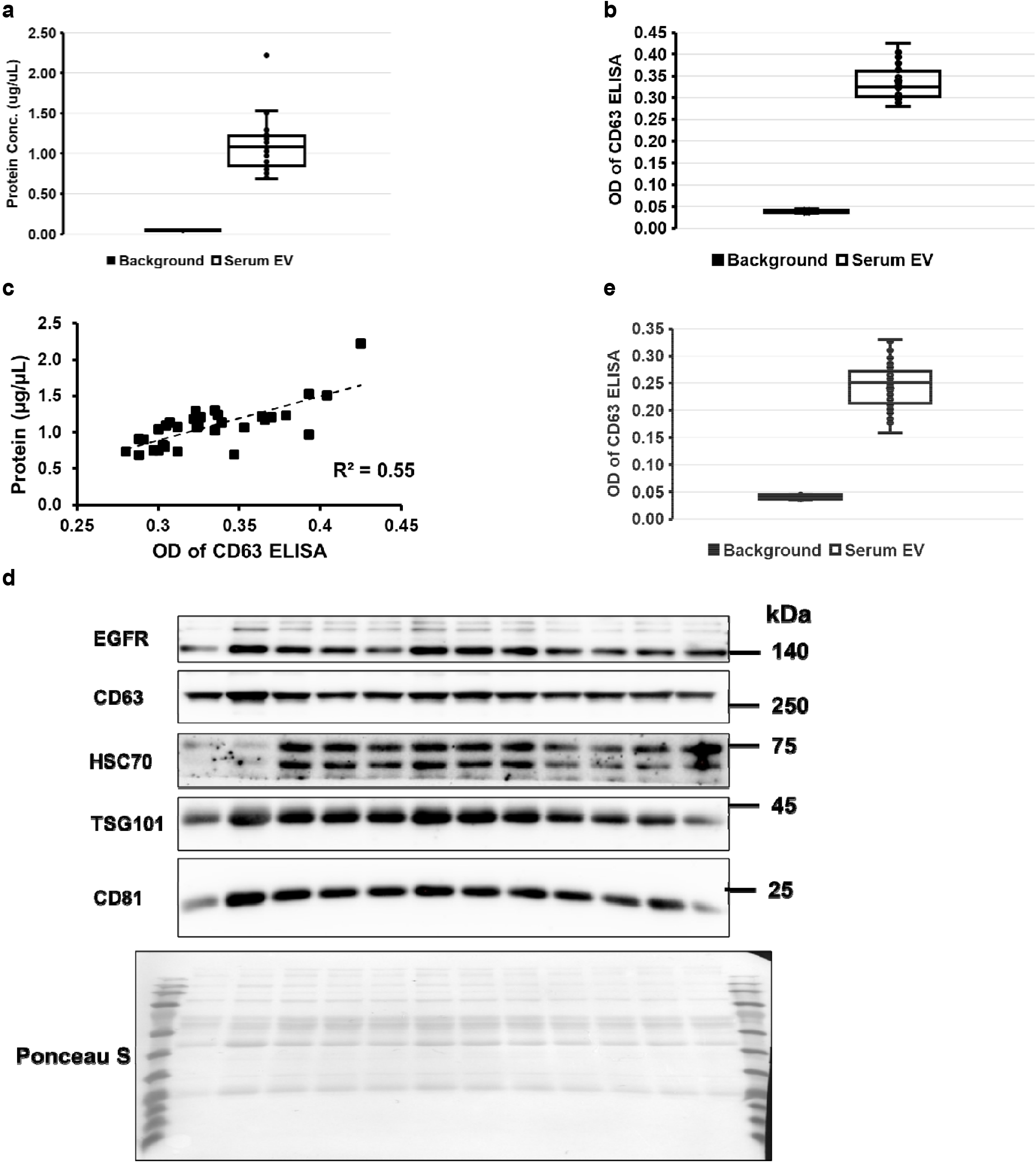
Characterization of EV isolated by MagPEG high throughput protocol. **(a)** Protein quantification of MagPEG purified EV in manual high throughput protocol. **(b)** CD63 direct ELISA read of the same sample as in (a). **(c)** Scatter plot of protein concentration and CD63 OD from the same sample as in (a) and (b). **(d)** Western blot for EV markers of 12 random samples from (a). **(e)** CD63 direct ELISA read of 96 samples from MagPEG automated high throughput protocol.

With the help from ICBR at University of Florida, we tested the MagPEG protocol with Mosquito HV Genomics and Dragonfly Discovery liquid handling platforms. As little as 5 µL serum was successfully used as input for serum EV isolation. CD63 direct ELISA results prove that similar high quality EV as from manual protocol (Fig.7b, e). Fully automated process also greatly reduces the handling time in isolation protocol. Total process time for 96 samples were only 45 minutes.

Summarized all the data we present, MagPEG has ability to isolate EVs on high throughput scale with low-cost, easy handling, high purity and time-saving features. Based on our knowledge, there is no current available method or product has similar capacity and performance.

### 3.5 MagPEG integrates extended protocol for EV associated nucleic acid and protein purification

For decades, magnetic beads and PEG combination has been considered as a gold standard for nucleic acid purification [9], and widely used in sequencing library preparation [17, 18]. In the MagPEG workflow, we chose the most common carboxyl modified magnetic beads, which is widely used in DNA/RNA isolation protocol [9, 18, 27]. To integrate its nucleic acid purification feature into our MagPEG workflow, we select the point that after the second wash (Fig.1a, Fig.10). On this step, EV still bond with beads, and only little buffer residue which allow us to introduce lysis / DNA binding buffer. The role of same beads in EV isolation were changed to DNA purification after this step. After lysis EV with 0.5% Triton, PEG and NaCl were increased to working conc. of 13% and 1.6M respectively, to facilitate DNA binding as the most commonly used protocol [9, 17, 18]. The result showed that this extended protocol purified high yield dsDNA with very low protein impurity (Fig.8a). Although using the same concentration of PEG (13%) and NaCl (1.6 M) as in cfDNA purification protocol, the dsDNA profile was quite different (Fig.8b). In the cfDNA protocol, Triton lysis all the membrane vesicles and release DNA into solution, and the chromatogram showed both the short (mono-, di-tri-nucleosomes) and long fragment of DNAs [28-31]. MagPEG were using a low PEG condition (4% - 6%) for EV binding before lysis, which will only allow high molecular weight DNA binding to the beads. The size selection capacity of PEG concentration has been applied in NGS library preparation for many years [17, 18, 32].

**Fig8.**
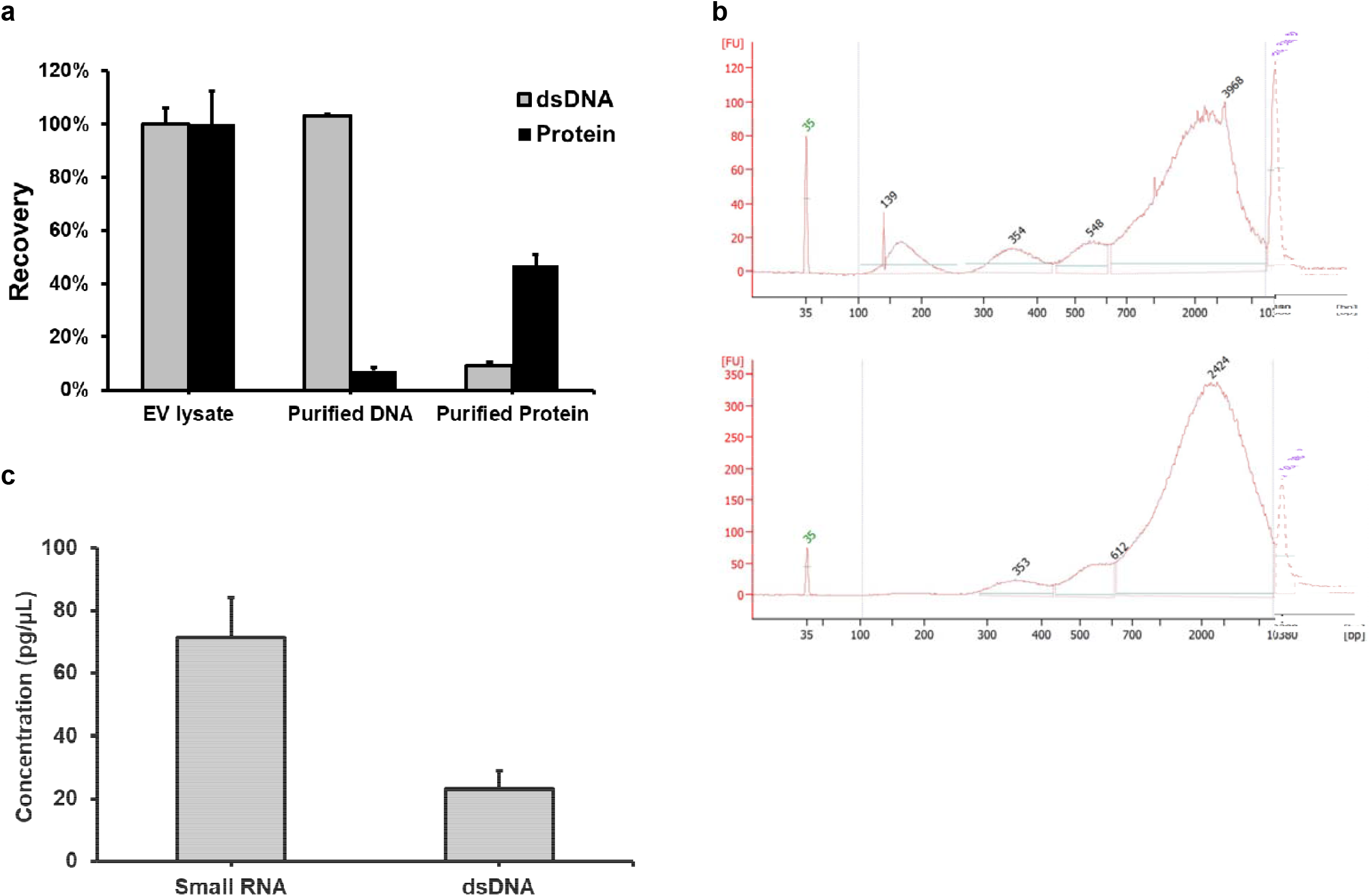
MagPEG extend protocol for protein / DNA / small RNA isolation. **(a)** Recovery of protein or dsDNA from MagPEG extend protocol. Samples were shown in triplicates. **(b)** Bioanalyzer profile of isolated dsDNA from serum (upper) or MagPEG EVs (lower). **(c)** Small RNA and dsDNA quantification results from the same sample purified by small RNA isolation protocol.

From protein isolation aspect, SP3 protocol had been used in our above result of proteomic sample preparation (Fig.6), but the beads were newly added after EV isolation. Here, we started also from the EV bond beads point to reuse the beads. In SP3 protocol, protein aggregate with beads is induced by 50% ethanol, which will also promote DNA binding. So, we added DNase to remove DNA in the lysis step, and then added ethanol to reach working concentration of 50%. After elution, the protein yield is about 50%, but with only 10% DNA left (Fig.8a). The difference between each MagPEG protocols were summarized in Table 5.

**Table 5.** MagPEG protocol comparison for different application

| Application | Pre-treatment | Binding | Wash | Elution |
| --- | --- | --- | --- | --- |
| EV isolation | - | PEG / NaCl<br>Low conc. | PBS<br>on magnet | PBS |
| DNA isolation | Lysis buffer<br>(0.5% Triton) | PEG / NaCl<br>High conc. | 70% Ethanol<br>on magnet | TE or H<sub>2</sub>O |
| Protein isolation | RDD buffer<br>(0.5% Triton)<br>DNase1 | Ethanol 50% | 80% Ethanol<br>off magnet | PBS<br>(SDS) |

Small RNA, especially microRNAs, is one of the important cargos carried by EVs [33, 34]. More and more researchers need to purify and characterize EV associate microRNAs. We also tested one of the common small RNA isolation protocols with magnetic beads. In this protocol, beads were first used to deplete large DNA/RNA under 20% PEG / 1.25M NaCl and discard. The supernatant incubated with new beads and promote small RNA binding in the present of 75% ethanol. The result showed that small RNA could be purified with low dsDNA contaminate (Fig.8c). We assumed adding DNase digestion step will further decrease dsDNA level in the final product. From these extended protocols, the magnetic beads in MagPEG could be reused to purify DNA, protein or small RNAs.

### 3.6 Covalent conjugation with beads allows multiplex protein profiling from MagPEG

Another interesting application of this carboxyl bead is it has the ability to covalent conjugate with protein or peptide. With the mediator of EDAC, carboxyl group will first be activated into o-acylisourea ester and then interact with primary amine group on protein N-terminus to form a covalent peptide bond [35]. This reaction could be performed under mild condition like room temperature, and without hash acidic, basic or denature solvent. After the reaction, protein will keep its nature formation, epitope or antigen binding capacity, which make it useful to create antibody conjugated beads. Here, we used the standard protocol to conjugate EV proteins onto the surface of carboxyl magnetic beads. The protocol needs to incubate beads with protein solution in MES buffer for 15 minutes before adding EDAC, but from our MagPEG protocol, as in DNA/protein purification, we could start from the same EV binding to beads point (Fig.1, Fig.10). MES buffer was used directly to resuspended beads, which will also greatly increase the conjugation efficiency (Supplementary Table 2). And the EDAC ratio was also been tested in our preliminary experiment to maximize protein conjugation without affect its detectability by ELISA. After the reaction was successfully done, PBST was used to remove by-product and other chemicals. Flow cytometry test of the conjugated beads shows some degree of aggregation after EDAC reaction in the 4% / 1.0M groups (Supplementary Fig.3).

Next, we tested the protein conjugated beads with standard ELISA protocol. Antibodies of three EV markers (CD63, CD9, CD81) were incubated separately with two MagPEG conditioned EV beads (6% / 0.2M and 4% / 1.0M), BSA conjugated beads with same protein amount and empty beads. The results (Fig.9a) showed that 6% / 0.2M had the stronger signal comparing with 4% / 1.0M sample, which is consistent with our previous western blot results (Fig.1h, Fig.5). And the background from control beads was all neglectable comparing with the sample signal.

Another protein profiling method we have previously described is TotalSeq assay [19]. The assay is using commercially available DNA conjugated antibody to quantify target proteins. As the EV protein had been immobilized on carboxy beads, we incubated the beads with TotalSeq antibody pool under the same protocol as in beads ELISA. Then, antibody conjugated DNA oligos were ligated with universal 3’-adaptor. The extension product can be quantified by qPCR with forward primer matched barcode region on DNA oligo and reverse universal primer on 3’-adaptor. As in beads ELISA experiment, BSA conjugated beads were also used as control. CD9 and CD81 did not show any positive signal on the BSA beads (Fig.9b), because the value of Ct_(control Ig)_ – Ct_(CD9)_ and Ct_(control Ig)_ – Ct_(CD81)_ was negative which means CD9/CD81 TotalSeq antibody has the same affinity as control IgG on beads. But EV protein conjugated beads presented positive signal, except CD63 antibody (Fig.9c). We assume that the clone of TotalSeq CD63 antibody (H5C6) recognize the N-terminus of CD63 which has been linked to carboxy beads, but the other CD63 antibody (TS63) we used in the beads ELISA has a different binding epitope which is not blocked in the EDAC reaction.

**Fig9.**
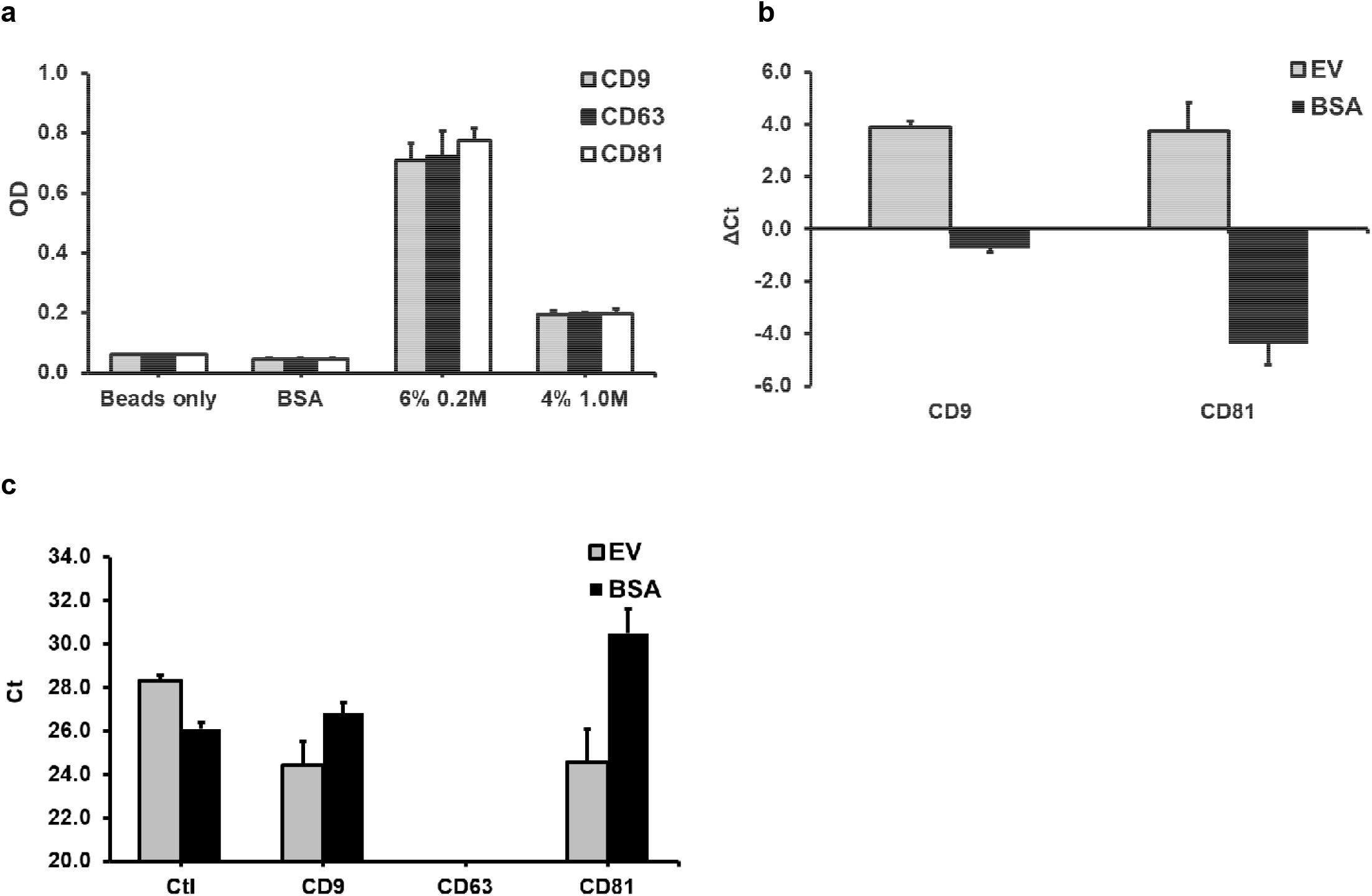
Covalent conjugation with beads for protein profiling by direct bead ELISA or TotalSeq assay. **(a)** Direct ELISA reads from CD9 / CD63 / CD81 antibody of beads, BSA conjugated beads, MagPEG 6% 0.2M or MagPEG 4 1.0M beads. **(b)** ΔCt (control Ig – CD9/CD81) value from BSA conjugated beads or MagPEG 6% 0.2M beads. **(c)** Ct value from BSA conjugated beads or MagPEG 6% 0.2M beads.

**Fig10.**
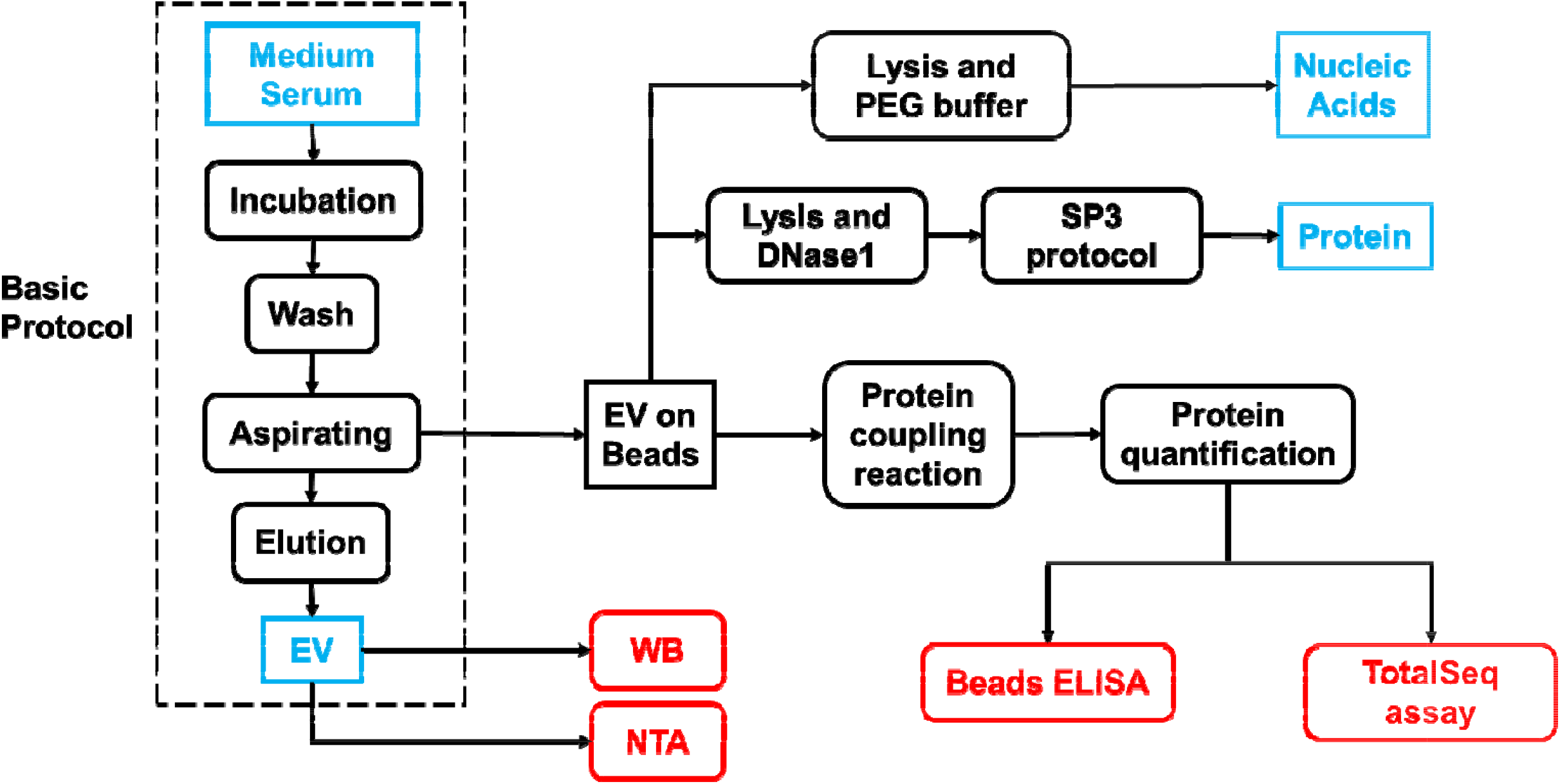
Complete MagPEG Protocol workflow. Input and output were shown in turquoise box, compatible assays were shown in red box.

By using MagPEG extended protocol of beads ELISA or TotalSeq assay, the whole experiment process from EV isolation to protein profiling could be completed in 8 hours with less labor- and time-intensive work than any other methods.

## 4. Discussion

For decades, PEG induced aggregation / precipitation had been widely applied in protein, nucleic acid, virus and EV isolations [4, 9, 10, 36, 37]. In this study, we merged ExtraPEG and magnetic beads to isolate high purity EVs in a fast, simply and easy way (Fig.10). A recent publication also showed that marker difference of CD63/CD81/CD9 indicate different EV subpopulations, like exosome and microvesicles [26]. There is no gold standard method to separate exosome from microvesicles, because their size, density and surface marker are always overlapped [38, 39]. Comparing with our previous ExtraPEG protocol, MagPEG purified EV subpopulations in different ratio under different conditions, which allowed user to adjust their own experiment conditions (Fig.1h, Fig.4, Fig.5). From the western blot and proteomic results, we believed that 6% / 0.2M condition enriched the most widely present EV populations include exosome, microvesicle and LDLs (Fig.1, Fig.4, Fig.5). Lipidomic study may be performed to further elucidate the question of LDL content. HSC70 had been observed absent in our MagPEG purified EVs under either condition, which is also shown in the results of ExtraPEG method [10]. Here, we make the same conclusion that HSC70 may belongs to a distinct EV subpopulation, or is just co-isolated under other methods.

PEG induced EV or cell fusion had been reported in several publications [40-42]. In our MagPEG protocol, induced EV fusion was also observed in 6% PEG group under longer incubation time (Fig.1e). Comparing with the overnight incubation in ExtraPEG protocol, shorter incubation time is a great improvement for EV isolation, which not only simplify the workflow but also lead to less protein co-precipitation and reduction of EV fusion or loss of biological function (Fig.1d-h).

In the MagPEG protocol incubation step, free DNA/RNA molecules (or proteins) were highly likely to be co-precipitated with EVs under PEG precipitation. Like the ultracentrifugation step in ExtraPEG, PBS wash step may also remove some of those outside associate nucleic acids, but the efficiency had not been tested. In the extended protocol for nucleic acid and protein purification, the EV binding beads were reused in the following step. To avoid carry over of co-precipitated nucleic acid or protein, eluted EV can be transferred into a new tube and mixed with new beads for protein and DNA isolation under the same protocol.

Fully automated or high throughput DNA isolation or purification method based on magnetic beads have been applied in academy or industry field for a long time [9]. There are several commercially available systems and kits from different suppliers (ThermoFisher, KingFisher Flex, MagMAX kit; Beckman, Biomek). Adapting from similar liquid handling platform and protocol, high throughput ability of MagPEG protocol had been proved.

For the EDAC coupling reaction between carboxyl beads and EV protein, we suggest to start on the EV bond beads step, which will greatly increase the conjugation efficiency (Supplementary Table 2). But similar to the concerns in the DNA/protein isolation protocol, to avoid carry over, EV can be transferred into a new tube and set up the EDAC reaction with original protocol. Another concern during the EDAC reaction is we do not know if membrane structure of EV had been disrupted. The only conclusion we know is this reaction keep the nature epitope form of EV protein which could be detect with antibody-based method (Fig.9). And theoretically, beads immobilized protein could also be analyzed by flow cytometry [6, 43] which has not been fully tested. On the aspect of TotalSeq assay, non-specific interactions of antibody-oligonucleotide conjugates with living cells had been reported recently [44]. This should be one of the reasons we could not get clear background in TotalSeq assay (Fig.9c) [19].

The recent EV isolation methodology studies are more focus on complex combinations or depend on novel instruments [45, 46], but we believe that “everything must be made as simple as possible”. Our MagPEG is a unique combination which is extended from well-developed techniques. Its simplicity and extendibility will greatly overcome the disadvantage of purity. The international Society for EV (ISEV) has emphasized the urgent need for more effective methods to isolate EV [1], and we believe that this procedure can fulfill that void. From the similarity of EV and virus, especially enveloped virus, we believe that MagPEG protocol will also succeed on virus isolation and characterization.

Take a look back on the whole picture of MagPEG protocol (Fig.10), we believe it has great potential to be applied in many fields and applications.

## Supporting information

Supplementary Proteomic data

## Author contributions

L.S. and D.G.M. conceived and designed the study. L.S., Y.Z. and D.G.M designed experiments and interpreted results. L.S., S.Y. and B.J. performed experiments. The manuscript was written by L.S. and D.G.M.. D.G.M. obtained funding to support the study.

