## Supplementary Figures for "MagPEG: a complete extracellular vesicle isolation/analysis solution"

**Supplementary Table 1. NTA result of elution from empty beads.**

| **Sample** | **Mean**  **(nm)** | **Mode**  **(nm)** | **SD** | **Conc.**  **(/uL)** |
| --- | --- | --- | --- | --- |
| PBS | 160 | 155 | 46 | 1.75E+08 |
| BSA-Beads | 147 | 137 | 26 | 4.40E+07 |

**Supplementary Table 2. Conjugated protein under different binding and EDAC condition**

|  | Conjugated total protein (ng) | | | |
| --- | --- | --- | --- | --- |
| EDAC ratio  Binding buffer | 1:1 | | 5:1 | |
| Ethanol (50%) | 1.3 | 1.3 | 1.7 | 2.3 |
| PEG (12%) | 1.3 | 0.9 | 1.0 | 1.3 |
| None | 0.8 | 0.6 | 0.5 | 0.8 |

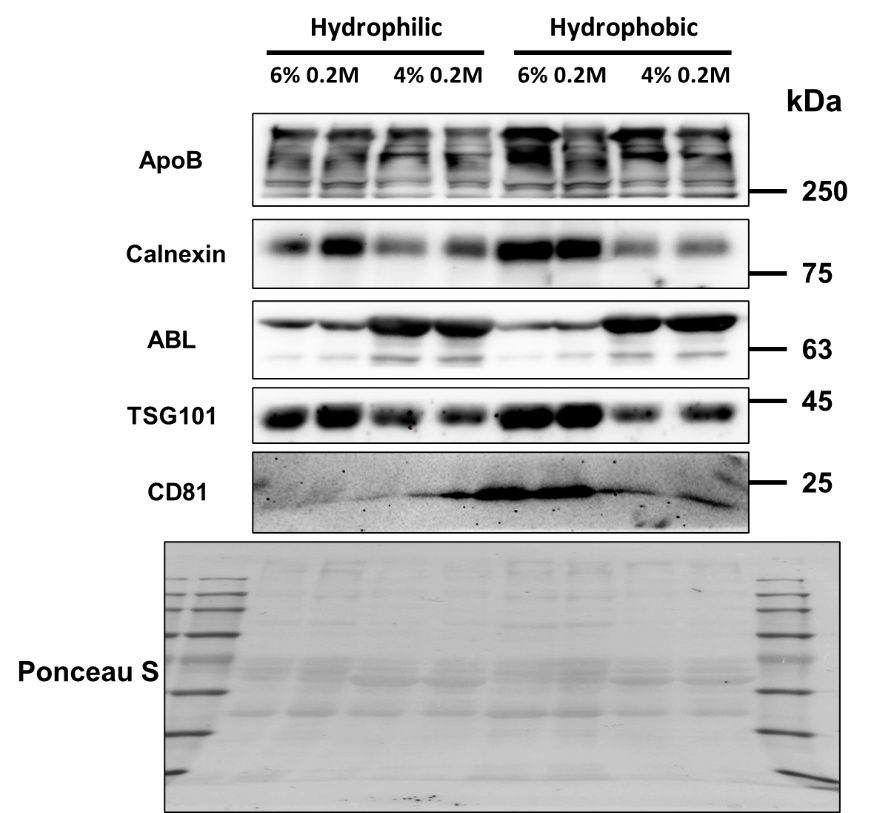

**Supplementary Fig 1. Western blot analysis of MagPEG protocol with different magnetic beads.** Duplicate samples were run for each condition.

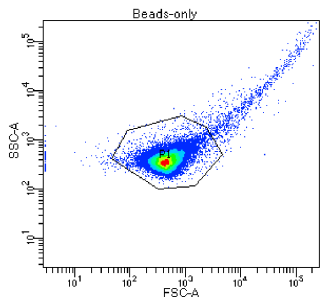

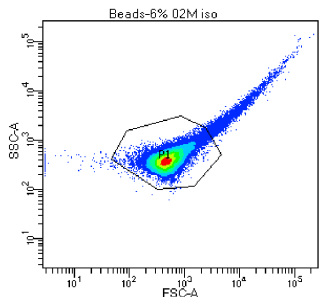

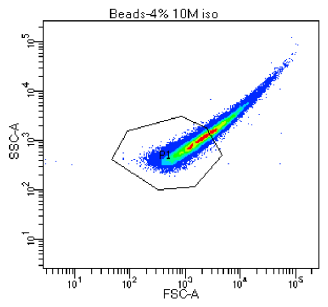

**Supplementary Fig 2. Flow cytometry analysis of protein conjugated beads.** Magnetic beads (left), MagPEG 6% 0.2M conjugated beads (middle) and MagPEG 4% 1.0M conjugated beads (right) were run under the same setting.
